# Neural correlates of object-extracted relative clause processing across English and Chinese

**DOI:** 10.1101/2022.09.12.507571

**Authors:** Donald Dunagan, Miloš Stanojević, Maximin Coavoux, Shulin Zhang, Shohini Bhattasali, Jixing Li, Jonathan Brennan, John Hale

## Abstract

Are the brain bases of language comprehension the same across all human languages, or do these bases vary in a way that corresponds to differences in linguistic typology? English and Mandarin Chinese attest such a typological difference in the domain of relative clauses. Using fMRI with English and Chinese participants, who listened to the same translation-equivalent story, we analyzed neuroimages time-aligned to object-extracted relative clauses in both languages. In a GLM analysis of these naturalistic data, comprehension was selectively associated with increased hemodynamic activity in left posterior temporal lobe, angular gyrus, inferior frontal gyrus, precuneus, and posterior cingulate cortex in both languages. This result suggests the processing of object-extracted relative clauses is subserved by a common collection of brain regions, regardless of typology. However, there were also regions that were activated uniquely in our Chinese participants albeit not to a significantly greater degree. These were in the temporal lobe. These Chinese-specific results could reflect structural ambiguity-resolution work that must be done in Chinese but not English ORCs.

## Introduction

To what degree does the brain basis of language processing vary across languages? (Bornkessel-Schlesewsky & Schlesewsky, 2016)? There is evidence in support of both variation (Wei et al., 2023) and universality (Dunagan et al., 2022; Malik-Moraleda et al., 2022). One approach to this question starts by identifying dimensions of typological difference between languages. This kind of investigation proceeds by comparing brain responses to linguistic expression types that manifest the given typological difference. The relative clause typology of English and Mandarin Chinese presents just such an opportunity. Within both psycholinguistics and neurolinguistics, these constructions have long been used to set-up controlled manipulations of language processing difficulty. Do they do so in a naturalistic comprehension scenario as well? The present article extends the neurobiological study of object-extracted relative clause processing across languages and into the naturalistic domain of audiobook listening (Hasson & Honey, 2012; Nastase et al., 2020; Willems, 2015). It uses publicly available fMRI data (J. T. Hale et al., 2022; J. Li et al., 2022; J. Li et al., 2021) to argue that indeed the language network is largely, but not entirely, uniform.

### Object-extracted relative clauses

A relative clause is a sentence-like grammatical unit that modifies a noun.

(1) [*_S_* you love the flower]
(2) the flower*_N_* that [*_S_* you love]

For instance, example 1 is an English sentence, a categorization that is indicated by square brackets labeled with the letter S. A variant of this simple sentence appears immediately to the right of the word “that” in the relative clause example 2. In this second example, the direct object “the flower” has gone missing from the S and surfaces instead as the head noun (N) of the entire expression. Quite generally, relative clauses (RCs) are sentence-like units that lack an element. The missing element, symbolized in example 2 with an underscore, receives its meaning from the head noun that the RC modifies (see Chaves & Putnam, 2021, for a more comprehensive overview). This meaning-related aspect is suggested by coindexation in examples 3 and 4 below, which comprise part of the materials used in the neuroimaging study reported here. These are all object-extracted RCs (henceforth: ORCs). This terminology reflects the fact that the missing element would have been the direct object rather than the subject of the relative clause verb – in this example,“love.” The words in square brackets in example 3 constitute the noun phrase that the ORC modifies. Its head noun (subscripted N in example 2) is referred to as the *filler*, while the position where that head noun makes its meaning-contribution is known as the *gap* (*site*). In English head nouns come before a relative clause; English RCs such as example 3 are postnominal. By contrast, Chinese RCs are prenominal. That is to say the relative clause occurs before the head noun which it modifies. This prenominal option, exercised by perhaps a quarter of the world’s languages (Dryer, 2013; see also Andrews, 1975; Downing, 1978), is exemplified in 4, where *DE* is a relativizing marker and *planet* is the filler. It is this typological difference in word order which makes the comparison of English and Chinese an intriguing prospect.

(3) [The flower]*_i_* that you love *_i_* is not in danger
(4) 小王子 来自 *_i_* 的 [星球]*_i_* 就 是 小 行星 B612 the little prince come from *_i_* DE planet*_i_* exactly is little planet B612 The planet that the little prince came from is asteroid B612

### Language processing and relative clauses

In English, results from reading time (Gibson et al., 2005; King & Just, 1991; Staub, 2010; Traxler et al., 2002; Traxler et al., 2005), question answering (Wanner & Maratsos, 1978), event related potential (ERP; King & Kutas, 1995; Müller et al., 1997), and neuroimaging (Caplan et al., 1985; Just et al., 1996; Stromswold et al., 1996) investigations converge on the conclusion that ORCs are more difficult for the human sentence processor than SRCs, thus making them a prime topic for investigation. In Chinese, however, things are not so clear. While some reading time (B. Chen et al., 2008; Hsiao & Gibson, 2003; Y. Lin & Garnsey, 2011; K. Xu et al., 2020a), ERP (Packard et al., 2011), and neuroimaging (K. Xu & Duann, 2020; K. Xu et al., 2020b) results indicate an ORC advantage (i.e., SRCs are more difficult to process than ORCs), other reading time (Jäger et al., 2015; C.-J. C. Lin & Bever, 2006, 2011; Vasishth et al., 2013), ERP (Xiong et al., 2019), and theoretical (Yun et al., 2015) results indicate a SRC advantage, which corresponds to the case in English. Further, some studies have found either no advantage (Lee & Chan, 2023; Zhou et al., 2018, fMRI and reading time, respectively), that the advantage is different at different points in the relative clause (Bulut et al., 2018, ERP), or that the advantage changes depending on whether the relative clause is subject- or object-modifying (Xiong & Newman, 2021, fMRI). A wide variety of theories have been put forward to account for (aspects of) this complex data pattern. The present study does not seek to decide among these theories, but rather leverages the fact that ORCs induce some sort of processing difficulty in English. While the literature is less unified about Chinese, ORCs appear to induce processing difficulty in that language as well, at least under some circumstances. With a view towards keeping as many factors as possible constant in a crosslinguistic comparison, the present study considers only ORCs.

Beyond their instrumental role in eliciting comprehension difficulty, ORCs are a valuable test case because of their size. Ranging in length from just a couple morphemes on up, ORCs occupy a middle ground between “small” and “large” linguistic expressions. This scale has not been previously investigated from a neuro-typological perspective, although neighboring scales have been. On the large end, previous work has found similar brain activations across Russian and English, as speakers listen to a translation-equivalent narrative (Honey et al., 2012). Patterns of activation in the default mode network can likewise identify a story, even when the representation was calculated from a translation of that story in another language (Dehghani et al., 2017). Indeed, one project using multisentence snippets from *Alice in Wonderland* has documented considerable cross-linguistic similarity across the language network (Malik-Moraleda et al., 2022).

By contrast, studies with smaller linguistic units have come to the opposite conclusion. Work on phonological access in Italian vs. English (Paulesu et al., 2000), pitch contour processing in Chinese vs. English (Gandour et al., 2003), and noun and verb representation in Chinese vs. Indo-European (P. Li et al., 2004) has all pointed to differences across languages. Examining units of intermediate size, such as ORCs, should help to fill-in this inconsistent picture.

### Current understanding of the brain basis

Previous neuroimaging work in English has associated ORC processing with activation in the left inferior frontal gyrus (IFG; Caplan et al., 2008; Just et al., 1996; Stromswold et al., 1996), right IFG (Just et al., 1996), left (posterior) middle/superior temporal gyrus ((p)M/STG; Caplan et al., 2008; Just et al., 1996), right pM/STG (Just et al., 1996), and left precuneus (Caplan et al., 2008). Previous neuroimaging work in Chinese has associated ORC processing with activation in the left IFG (Lee & Chan, 2023; Xiong & Newman, 2021), right IFG (Xiong & Newman, 2021), left pSTG (K. Xu & Duann, 2020; K. Xu et al., 2020b), right pSTG(K. Xu et al., 2020b), left mid M/STG (Lee & Chan, 2023), right mid M/STG (Lee & Chan, 2023), left anterior MTG (Xiong & Newman, 2021; K. Xu & Duann, 2020; K. Xu et al., 2020b), right anterior MTG (K. Xu et al., 2020b), left MTG broadly (Xiong & Newman, 2021), right MTG broadly (Xiong & Newman, 2021), left angular gyrus (AG; Xiong & Newman, 2021), right AG (Xiong & Newman, 2021), left premotor cortex (K. Xu & Duann, 2020; K. Xu et al., 2020b), right premotor cortex (Lee & Chan, 2023), left precuneus (Xiong & Newman, 2021), left medial frontal gyrus (Xiong & Newman, 2021), left fusiform gyrus (K. Xu & Duann, 2020; K. Xu et al., 2020b), left posterior cingulate cortex (K. Xu & Duann, 2020; K. Xu et al., 2020b), left temporal pole (Xiong & Newman, 2021), and right temporal pole (Xiong & Newman, 2021).

### Naturalistic language comprehension

Our current understanding of the brain basis of ORC processing is largely founded on controlled studies using decontextualized linguistic stimuli. How well do those conclusions generalize to everyday language comprehension? Proponents of naturalistic stimuli argue that narratives are a step in the right direction. For instance Hasson et al. (2010), in review, find that naturalistic stimuli, including audiobooks, can evoke more reliable and functionally-selective responses than experimental stimuli. As well, in an investigation of the neural correlates of code switching in bilinguals with MEG, Blanco-Elorrieta and Pylkkänen (2017) find that production and comprehension results vary dramatically depending on whether the stimuli are artificial or naturalistic. The key difference between the present study and earlier work with non-naturalistic stimuli is an enriched discourse context, one that one that “grounds” storybook characters in a storybook world (Hasson et al., 2018).

### Open questions and hypotheses

The considerations mentioned above underline a tension that currently exists between neural uniformity and typological variability. The present study examines that tension in the domain of ORCs. It tests whether the brain bases of ORC processing are the same for English and Chinese, languages where ORCs manifest a well-known distinction between prenominal and postnominal word order. It is hypothesized that: (1) the correlates for processing ORCs will be the same across the two languages and implicate at least the left inferior frontal gyrus and left posterior temporal lobe; and (2) that the naturalistic stimuli will evoke activation in brain areas outside of the traditional language network. To test these hypotheses, we analyzed ORCs in naturalistic fMRI data collected while English and Chinese participants listened to translation-equivalent audiobooks of the *Le Petit Prince* (*The Little Prince*), by de Saint-Exupéry, as detailed below.

## Materials and methods

The fMRI data analyzed were *The Little Prince Datasets* (J. T. Hale et al., 2022; J. Li et al., 2022; J. Li et al., 2021), a published collection of datasets in which Chinese and English participants were scanned while they engaged in the naturalistic process of listening to a translation-equivalent audiobook in their native language. The participants listened to the children’s story *The Little Prince*. A general linear model (GLM) analysis was performed which included a binary regressor for ORC processing as well as control regressors of non-interest. The binary ORC regressor marks with a 1 only those words where ORC processing occurs. For this binary ORC metric, all other words are marked as 0. This can be visualized in Fig. 1, and more detail will be given below. In order to ensure the fidelity of any effects found for the ORC metric, control regressors were included in the first level of the GLM for speaker amplitude, speaker pitch, spoken word rate, word frequency, syntactic processing, word-by-word surprisal, and lexical semantics. In this way, any effects found for the ORC metric cannot be attributed to these alternate aspects of language processing. The first-level English and Chinese ORC brain maps were entered into a second-level GLM analysis with a two-sample t-test design matrix encoding. Effects of ORC processing in each language were analyzed as was the difference in ORC processing between the two languages. Further, to probe cross-linguistic commonality between the two languages, the voxel-wise intersection was calculated between the ORC processing effects in English and Chinese. That is, if a voxel was found to be activated by ORC processing for both English and Chinese participants, it is recorded in order to construct a new brain map indicating voxels which are commonly activated during ORC processing for both English and Chinese participants.

**Figure 1.**
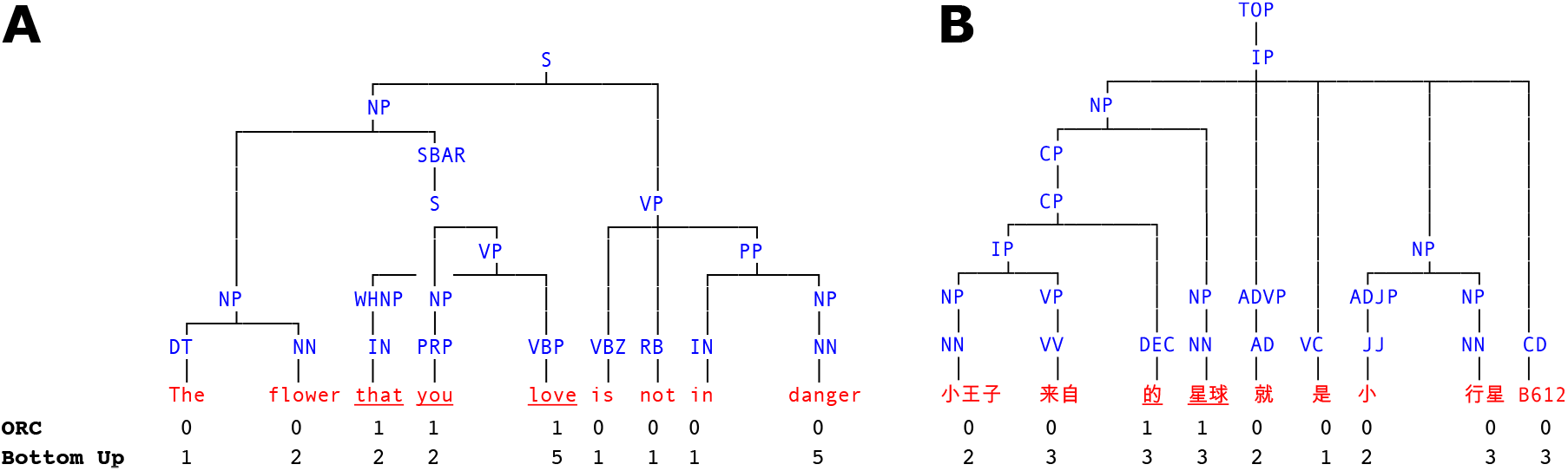
Example phrase structure trees with annotations for the corresponding word-by-word object-extracted relative clause (ORC) and bottom up syntactic processing metric values. A, The example English sentence with an object-extracted relative clause given in example 3. B, The example Chinese sentence with an object-extracted relative clause given in example 4.

### fMRI data

#### Participants

Chinese participants were 35 healthy, right-handed young adults (15 females, mean age=19.3, SD=1.6). They self-identified as native Chinese speakers, and had no history of psychiatric, neurological, or other medical illness that could compromise cognitive functions. All participants were paid, and gave written informed consent prior to participation, in accordance with the IRB guidelines of Jiangsu Normal University.

English participants were 49 young adults (30 females, mean age=21.3, SD=3.6) with no history of psychiatric, neurological or other medical illness that might compromise cognitive functions. They self-identified as native English speakers and strictly qualified as right-handed on the Edinburgh handedness inventory. All participants were paid, and gave written informed consent prior to participation, in accordance with the IRB guidelines of Cornell University.

#### Procedure

After giving their informed consent, participants were familiarized with the MRI facility and assumed a supine position on the scanner. They were instructed to not move as best as they could throughout scanning as movement would make the scans unusable. Next, participants were put in the head-coil with pillows under and on the sides of their head and under the knees for comfort and to reduce movement over the scanning session. Participants were given a bulb in their right hand and told to squeeze if something was wrong or they needed a break during scanning. Once in place, participants chose an optimal stimulus volume by determining a level that was loud but comfortable. Auditory stimuli were delivered through MRI-safe, high-fidelity headphones inside the head coil (English: Confon HP-VS01, MR Confon, Magdeburg, Germany; Chinese: Ear Bud Headset, Resonance Technology, Inc, California, USA). The headphones were secured against the plastic frame of the coil using foam blocks.

The English and Chinese participants went through one scanning session, which was divided into 9 runs, with each lasting for about 10 minutes. Participants listened passively to 1 section of the audiobook in each run and completed 4 quiz questions after each run (36 questions in total). These questions were used to confirm their comprehension and were viewed by the participants via a mirror attached to the head coil and they answered through a button box. During scanning, participants were monitored by a camera over their left eye. If they appeared drowsy or seemed to move too much, the operator of the scanner gave them a warning over the intercom by producing a beep or speaking to them. During breaks between the runs, participants were told that they could relax but not move. Finally, participants were paid and sent home. The entire session lasted for around 2.5 hours.

#### Acquisition

MRI images were acquired with a 3T MRI GE Discovery MR750 scanner with a 32-channel head coil. Anatomical scans were acquired using a T1-weighted volumetric Magnetization Prepared Rapid Gradient-Echo pulse sequence. Functional scans were acquired using a multi-echo planar imaging sequence with online reconstruction (TR=2000 ms; TEs=12.8, 27.5, 43 ms; FA=77*^◦^*; matrix size=72 × 72; FOV=240.0 mm × 240.0 mm; 2× image acceleration; 33 axial slices; voxel size=3.75 × 3.75 × 3.8 mm).

#### Preprocessing

MRI data files were converted from DICOM to NIfTI format and preprocessed using AFNI (Cox, 1996, version 16). *Anatomical*. The anatomical/structural MRI scans were deskulled using *3dSkullStrip*. The resulting anatomical images were nonlinearly aligned to the Montreal Neurological Institute N27 template brain. Resulting anatomical images were used to create grey matter masks. *Functional*. The first 4 volumes in each run were excluded from analyses to allow for T1-equilibration effects. The fMRI timeseries were then corrected for slice-timing differences (*3dTshift*) and despiked (*3dDespike*). Next, volume registration was done by aligning each timepoint to the mean functional image of the center timeseries (*3dvolreg*). Then the volume-registered and anatomically-aligned functional data were nonlinearly aligned to the MNI template brain. Multi-echo independent components analysis (Kundu et al., 2012) was used to denoise data for motion, physiology, and scanner artifacts. Images were then resampled at 2 mm cubic voxels (*3dresample*).

### Stimuli and storybook annotations

Participants listened to a translation-equivalent audiobook of *The Little Prince* in its entirety. The English audiobook is 94 minutes long, translated by David Wilkinson, and read by Karen Savage. The Chinese audiobook is 99 minutes long and read by a professional female Chinese broadcaster hired by the experimenter.

Each word in the storybooks was annotated for a number of metrics which were expected to be cognitively influential. The first step in annotating the storybook texts was to parse them for syntactic structure. The syntactic annotations, or trees, were used to calculate a word-by-word syntactic complexity metric and to calculate a binary word-by-word ORC annotation. The English text was parsed using the mtgpy parser (Coavoux, 2021), while the Chinese text was parsed using the benepar parser (Kitaev et al., 2019). The performance of these systems is given in the supplementary material. Following J. Brennan et al. (2012), a word-by-word “bottom-up” complexity metric corresponding to the count of reduce operations in an incremental shift-reduce parser was defined using the parse trees. J. Li and Hale (2019, §7.2.3.1) discuss the relationship between the complexity metric and the parsing strategy at greater length. This includes a worked example and a table of complexity values, analogous to the metric used here. This coregressor serves to control for effects of syntactic processing and can be visualized in Fig. 1. The use of 1-best parses means that the metric is blind to any sort of ambiguity resolution work.

Tree-geometric properties were used to find the ORC constructions. An example can be given in roughly the notation used by Richard Pito’s 1992 tgrep, tgrep2 (Rohde, 2005), and tregex (Levy & Andrew, 2006). Example 5 gives a tregex pattern for identifying ORC constructions in English. The notation *A < B* means that *B* is a child of *A* and *A << B* means that *B* is a descendant of *A*. This pattern metaphorically looks for a WH tag that is dominated by an NP that contains an SBAR with an object gap. The symbol SBAR identifies sentence-like units, including initial complementizers such as “that” which appears in example 2 on page 3. SBAR includes initial complementizers whereas S does not (see Marcus et al., 1993).

The pattern in 5 matches the structure in Fig. 1A.

(5) NP < (SBAR < (S < NP) < < /WH.*/)

The first instance of the symbol NP in pattern 5 matches the filler. To find gaps, a heuristic is used. This heuristic is based on the smallest sentence-like constituent which encloses the RC (variously SBAR, SQ, Srel, SENT or CP). Within that constituent, a gap is postulated immediately adjacent to the head verb. These are located via the head-finding rules of Collins (2003) and their equivalents for Chinese borrowed from CoreNLP (Manning et al., 2014). The tregex expression for identifying ORC constructions in Chinese is given in the supplementary material. The number of identified ORC constructions for each language is given in Table 1. Each observation was manually checked by native speakers who are also trained linguists.

**Table 1.** The number of identified English and Chinese object-extracted relative clauses in The Little Prince.

|  | Object-extracted relative clauses |
| --- | --- |
| English | 32 |
| Chinese | 30 |

Annotation of the binary ORC metric proceeded in correspondence to areas of expected increased cognitive demand. That is, words where increased cognitive demand associated with ORC processing was expected were annotated with a 1, while all other words were annotated with a 0. For English ORCs, each word between the filler and the gap site was annotated with a 1 (Wanner & Maratsos, 1978). For Chinese ORCs, each word between the gap site and the end of the filler was annotated with a 1. One may notice that in the English case, the area of expected increased processing demand corresponds to the ORC itself, but in the Chinese case, the area of expected increased demand corresponds to the filler rather than the ORC. As discussed in the Chinese relative clause processing literature (Bulut et al., 2018; Hsiao & Gibson, 2003; Jäger et al., 2015; C.-J. C. Lin & Bever, 2011; Vasishth et al., 2013), in Chinese, there is main clause/relative clause structural ambiguity during the actual relative clause which is not resolved until at least the relativizer *DE*. Following the relativizer, the filler element must be processed as the head noun which the ORC is modifying. Fig. 1A gives the parse tree for example 3. The underlined words, *that you love*, indicate the span between the filler and the gap site where an increase in cognitive demand is expected in English. Fig. 1B gives the tree for example 4. The underlined words, *DE planet*, indicate the span from the gap site through the filler and indicate where increased cognitive demand is expected in Chinese.

Additionally, pretrained autoregressive transformer language models—one for each language—were used to calculate word-by-word surprisal (J. Hale, 2001; Levy, 2008). Surprisal has been shown to be associated with activity in the language network (J. R. Brennan et al., 2016; Henderson et al., 2016), and was included to control for incremental sequential processing. Furthermore, word-by-word lexical surprisal is particularly pertinent to Chinese ORC constructions because, as just mentioned, Chinese relative clauses have main clause/relative clause structural ambiguity at least until the relativizer *DE*. Recent work (Oh & Schuler, 2022) has shown that as pretrained language model size increases, the ability to use surprisal to predict reading time decreases. For this reason, we use GPT2-caliber models. As measured by number of parameters, the two models are reasonably similar in complexity. For English, GPT2 (Radford et al., 2019, 1.5 billion parameters and trained on 40 gigabytes of data) was used. For Chinese, Chinese GPT2 (Zhao et al., 2019, 102 million parameters and trained on 14 gigabytes of data) was used. The Chinese model was accessed via HuggingFace Transformers (Wolf et al., 2019).

Lastly, as a lexical semantic control, word vectors were taken from the pretrained English and Chinese fastText models (Bojanowski et al., 2017; Grave et al., 2018). Word vectors are numerical representations which encode distributional properties of lexical items; these properties often reflect lexical meaning distinctions. The fastText word vectors were trained on Common Crawl and Wikipedia using Continuous Bag of Words with position-weights, in dimension 300, with character n-grams of length 5, and a window of size 5 and 10 negatives. Native from the model, the vector for each word is of dimensionality 300. That is, each word is associated with 300 real-valued numbers. In order utilize the word vectors in the GLM analysis, they are reduced down to 5 dimensions via principal component analysis with the included utility from the fastText toolkit.

## Experimental design and statistical analysis

### GLM analysis

The GLM analysis was carried out using Nilearn (Abraham et al., 2014; Pedregosa et al., 2011, version 0.9.1), a package for the statistical analysis of neuroimaging data in Python. Low-level regressors of non-interest included spoken word-rate, speaker pitch, and root mean squared amplitude of the spoken narration (RMS). These coregressors served to ensure that any results of interest were not the result of low-level language processing (cf. Bullmore et al., 1999; Lund et al., 2006). Word-by-word regressors of non-interest included log lexical frequency (log unigram frequency from the Google Books Ngram Viewer), the bottom-up syntactic processing metric, large language model surprisal, and word vectors. The regressor of interest was the word-by-word object relative metric.

All predictors were convolved with the SPM canonical hemodynamic response function. Following convolution, the lexical frequency, bottom-up syntactic processing, surprisal, word vector, and ORC word-by-word regressors were orthogonalized with respect to word-rate to remove correlations resulting from their common timing. Before modeling, the predictors were standardized (shifted to mean 0 and scaled to standard deviation 1) by scanning session/storybook section.

Throughout the GLM analysis, a liberal cortical mask (https://surfer.nmr.mgh. harvard.edu/fswiki/CorticalParcellation_Yeo2011) was applied, calculated from the 1,000 participants in Buckner et al. (2011) and Yeo et al. (2011). At the first level of the GLM, linear models were fit to the voxel blood oxygen level dependent (BOLD) timecourses. For the English participants, the English-associated regressors were used. For the Chinese participants, the Chinese-associated regressors were used. In all other respects, the first level models were the same. The Wilkinson-Rogers formula for both language-specific, first level GLMs is given in 6, where word_rate marks the offset of every word with a 1, RMS is root mean squared amplitude of the spoken narration and is marked every 10 ms, freq marks the offset of every word with its log lexical frequency, f0 is the speaker pitch and is marked every 10 ms, bottom_up marks the offset of every word with its syntactic complexity metric value, GPT2_surprisal marks the offset of every word with how surprised the language model is that it encounters the word given the preceding context, **word_vectors**_5_ marks the offset of every word with 5 regressors corresponding to the values of the word’s pretrained fastText vector, following a model reduction from word vector dimensionality 300 to dimensionality 5, and obj_relative marks the offset of every word with the binary ORC regressor value at that word.
(
(6) BOLD ∼ 1 + word_rate + rms + freq + f 0 + bottomup + GPT 2_surprisal + **word_vector**_5_ + obj_relative

The nine scanning sessions/storybook sections were used to compute fixed effects for each participant. Data for two Chinese participants resulted in errors at the first level and were not further analyzed, leaving data from 33 Chinese participants.

The first-level object relative coefficient maps were entered into the second level of the GLM analysis with a two-sample t-test design matrix encoding:

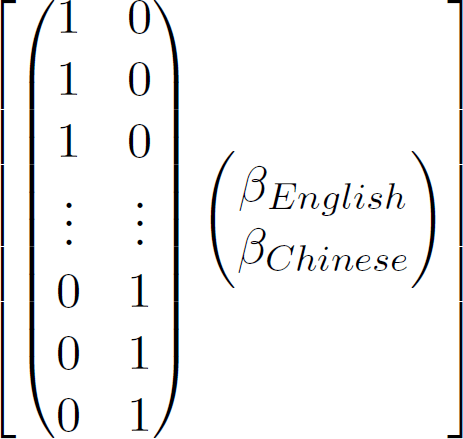

An equivalent number of English and Chinese first-level maps were analyzed: 33 of each language. An 8 mm full width at half maximum Gaussian smoothing kernel was applied to counteract inter-participant anatomical variation. Effects of object relative clause processing were calculated for English and Chinese, as were the contrasts between the two languages, that is, Chinese *>* English and English *>* Chinese. In this specific case of a two-sample t-test, these contrasts will result in the same voxel maps, but with flipped signs. Further, the intersection of voxels found to be significant in each of the two languages was calculated in order to identify common voxels associated with object relative processing.

The presented English and Chinese maps are z-valued and thresholded with an expected false discovery rate (FDR) *<* .05 and a cluster threshold of 125 voxels. The intersection map is the voxel-level intersection of the FDR-thresholded English and Chinese maps with no cluster thresholding. The MNI2TAL tool from the Yale BioImage Suite (Lacadie, Fulbright, et al., 2008; Lacadie, Fulbright, et al., 2008, version 1.4) was referenced for brain region and Brodmann area labels.

#### Data and code availability

The fMRI data are available through the associated OpenNeuro repository (https://openneuro.org/datasets/ds003643/versions/2.0.1). The analysis code is publicly available on GitHub, but the link has been retracted for anonymity during peer review.

## Results

### Behavioral results

Participants answered four four-choice comprehension questions after each section (36 questions in total). Participants performed well with a mean accuracy of 86.4% (SD=2.7) for Chinese participants and 89.5% (SD=3.8) for English participants.

### GLM

For the English participants, ORC processing was associated with an increase in activation in left pMTG, extending into the fusiform gyrus, left AG extending into the precuneus and posterior cingulate cortex, the left IFG, including the pars opercularis and pars triangularis, and left premotor cortex and superior frontal lobe. Decreases in activation were seen in left primary auditory cortex, bilateral precuneus, right anterior frontal lobe, and right premotor cortex. These increases and decreases in activation can be seen in Fig. 2 and more detail can be found in Table 2.

**Figure 2.**
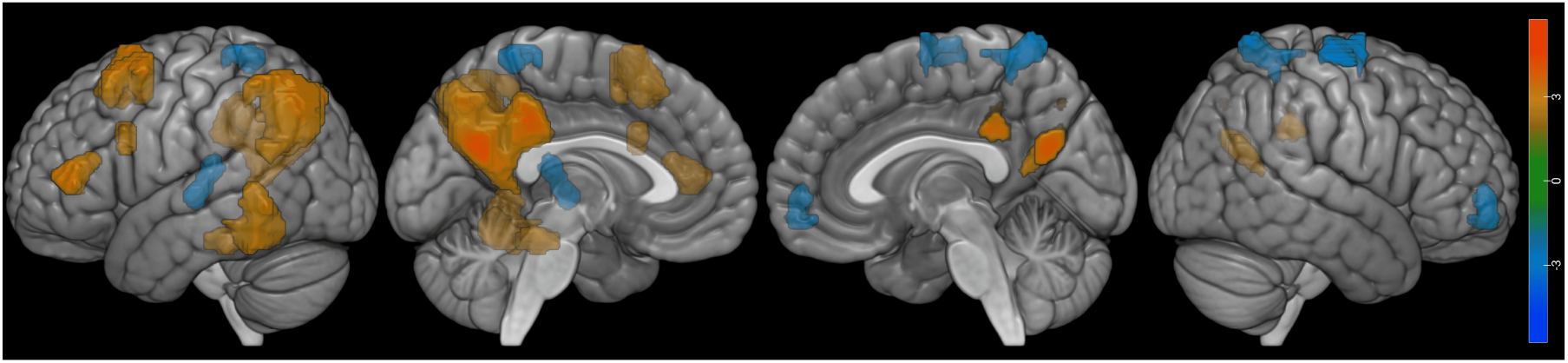
English object-extracted relative clause GLM results (z-valued), thresholded with an expected false discovery rate < .05 and a cluster threshold of 125 voxels.

**Table 2.** English object-extracted relative clause GLM results, thresholded with an expected false discovery rate < .05 and a cluster threshold of 125 voxels. Region, cluster size, MNI coordinates, and peak statistic (z).

| Region | Cluster size<br>(mm<sup>3</sup>) | MNI coordinates |  |  | Peak stat (z) |
| --- | --- | --- | --- | --- | --- |
|  |  | x | y | z |  |
| L Angular Gyrus (BA 39) | 35488 | -38.0 | -70.0 | 36.0 | 5.75 |
| L Posterior Cingulate Cortex (BA 31) |  | -8.0 | -62.0 | 24.0 | 5.75 |
| L Angular Gyrus (BA 39) |  | -36.0 | -58.0 | 38.0 | 5.24 |
| L Posterior Cingulate Cortex (BA 23) |  | -6.0 | -36.0 | 36.0 | 5.20 |
| L Premotor Cortex (BA 6) | 6504 | -30.0 | 14.0 | 58.0 | 4.90 |
|  |  | -24.0 | 14.0 | 68.0 | 4.63 |
| L Fusiform Gyrus (BA 37) | 2280 | -30.0 | -34.0 | -22.0 | 4.77 |
| L Fusiform Gyrus (BA 37) | 6584 | -52.0 | -48.0 | -20.0 | 4.75 |
| L Middle Temporal Gyrus (BA 21) |  | -58.0 | -50.0 | -6.0 | 3.86 |
| L Pars Triangularis (BA 45) | 2480 | -48.0 | 40.0 | 6.0 | 4.19 |
| L Pars Opercularis (BA 44) | 1136 | -36.0 | 14.0 | 30.0 | 3.43 |
|  |  | -48.0 | 14.0 | 20.0 | 3.04 |
| R Precuneus (BA 7) | 2832 | 16.0 | -56.0 | 72.0 | -4.57 |
|  |  | 18.0 | -40.0 | 66.0 | -3.41 |
|  |  | 16.0 | -44.0 | 56.0 | -3.36 |
|  |  | 14.0 | -34.0 | 66.0 | -3.05 |
| L Primary Auditory Cortex (BA 41) | 1880 | -34.0 | -26.0 | 12.0 | -4.04 |
|  |  | -40.0 | -20.0 | 2.0 | -3.94 |
| R Anterior Frontal (BA 10) | 1688 | 30.0 | 52.0 | -4.0 | -3.83 |
| L Precuneus (BA 7) | 1376 | -16.0 | -44.0 | 66.0 | -3.76 |
| R Premotor Cortex (BA 6) | 2200 | 22.0 | -4.0 | 74.0 | -3.55 |
|  |  | 34.0 | -14.0 | 72.0 | -3.55 |
|  |  | 44.0 | -10.0 | 64.0 | -3.44 |
|  |  | 26.0 | -4.0 | 70.0 | -3.40 |

For the Chinese participants, ORC processing was associated with an increase in activation in left pM/STG, bilateral mid and anterior STG, right temporal pole, left AG, left IFG, including the pars opercularis and pars triangularis, bilateral premotor cortex, bilateral posterior cingulate cortex, and bilateral precuneus. Decreases in activation were seen in the right occipital lobe and right fusiform gyrus. These increases and decreases in activation can be seen in Fig. 3 and more detail can be found in Table 3.

**Figure 3.**
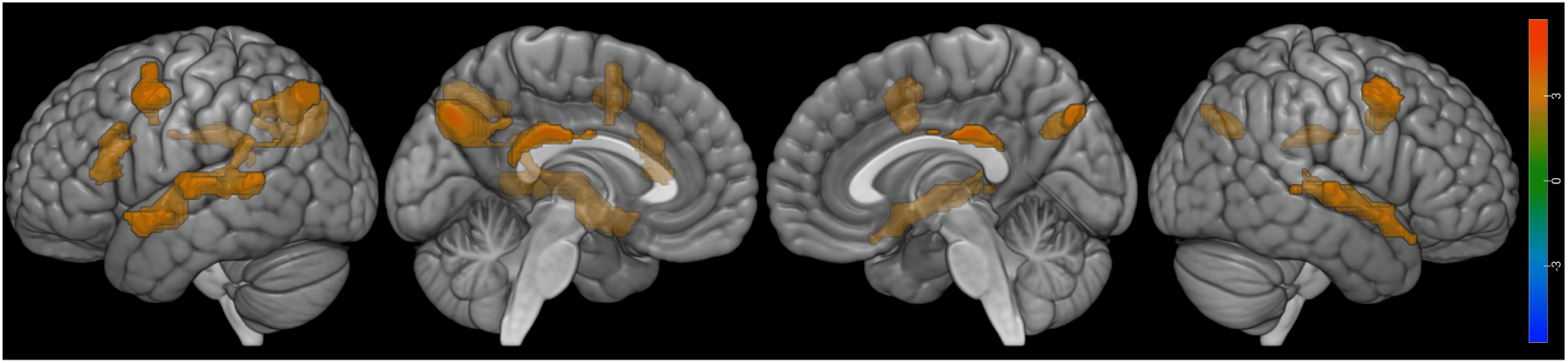
Chinese object-extracted relative clause GLM results (z-valued), thresholded with an expected false discovery rate < .05 and a cluster threshold of 125 voxels.

**Table 3.** Chinese object-extracted relative clause GLM results, thresholded with an expected false discovery rate < .05 and a cluster threshold of 125 voxels. Region, cluster size, MNI coordinates, and peak statistic (z).

| Region | Cluster size<br>(mm<sup>3</sup>) | MNI coordinates |  |  | Peak stat (z) |
| --- | --- | --- | --- | --- | --- |
|  |  | x | y | z |  |
| L Precuneus (BA 7) | 6960 | -8.0 | -72.0 | 34.0 | 5.69 |
| R Precuneus (BA 7) |  | 18.0 | -64.0 | 32.0 | 3.97 |
|  |  | 6.0 | -74.0 | 38.0 | 3.75 |
| L Posterior Cingulate Cortex (BA 23) |  | -10.0 | -50.0 | 34.0 | 3.19 |
| L Posterior Cingulate Cortex (BA 23) | 3128 | -6.0 | -34.0 | 24.0 | 5.36 |
| R Posterior Cingulate Cortex (BA 23) |  | 2.0 | -30.0 | 26.0 | 4.81 |
| L Premotor Cortex (BA 6) | 3560 | -52.0 | 4.0 | 48.0 | 4.90 |
|  |  | -40.0 | 2.0 | 56.0 | 3.70 |
| L Angular Gyrus (BA 39) | 3712 | -44.0 | -72.0 | 48.0 | 4.76 |
|  |  | -38.0 | -62.0 | 44.0 | 4.43 |
| R Premotor Cortex (BA 6) | 3544 | 52.0 | 10.0 | 48.0 | 4.49 |
| R Frontal (BA 8) |  | 34.0 | 4.0 | 36.0 | 4.10 |
|  |  | 42.0 | 8.0 | 38.0 | 4.04 |
| L Superior Temporal Gyrus (BA 22) | 10528 | -62.0 | -18.0 | 6.0 | 4.40 |
|  |  | -50.0 | -2.0 | -8.0 | 4.20 |
| L Middle Temporal Gyrus (BA 21) |  | -60.0 | -42.0 | 6.0 | 4.10 |
| L Angular Gyrus (BA 39) |  | -50.0 | -42.0 | 22.0 | 3.64 |
| R Superior Temporal Gyrus (BA 22) | 6392 | 54.0 | -4.0 | -4.0 | 4.27 |
|  |  | 56.0 | -24.0 | 4.0 | 3.77 |
| R Temporal Pole (BA 38) |  | 52.0 | 20.0 | -20.0 | 3.74 |
| R Superior Temporal Gyrus (BA 22) |  | 66.0 | -16.0 | 2.0 | 3.71 |
| L Pars Opercularis (BA 44) | 2696 | -46.0 | 22.0 | 18.0 | 3.93 |
|  |  | -34.0 | 22.0 | 24.0 | 3.89 |
| L Pars Triangularis (BA 45) |  | -44.0 | 20.0 | 12.0 | 3.81 |

Performing the English *>* Chinese and Chinese *>* English contrast analyses, no voxels survive expected FDR *<* .05 thresholding.

Commonly activated for both Chinese and English participants in response to ORCs were left pMTG, left angular gyrus, left IFG, including the pars opercularis and pars triangularis, left premotor cortex, and left posterior cingulate cortex and precuneus. The commonalities can be seen in Fig. 4 and more detail can be found in Table 4.

**Figure 4.**
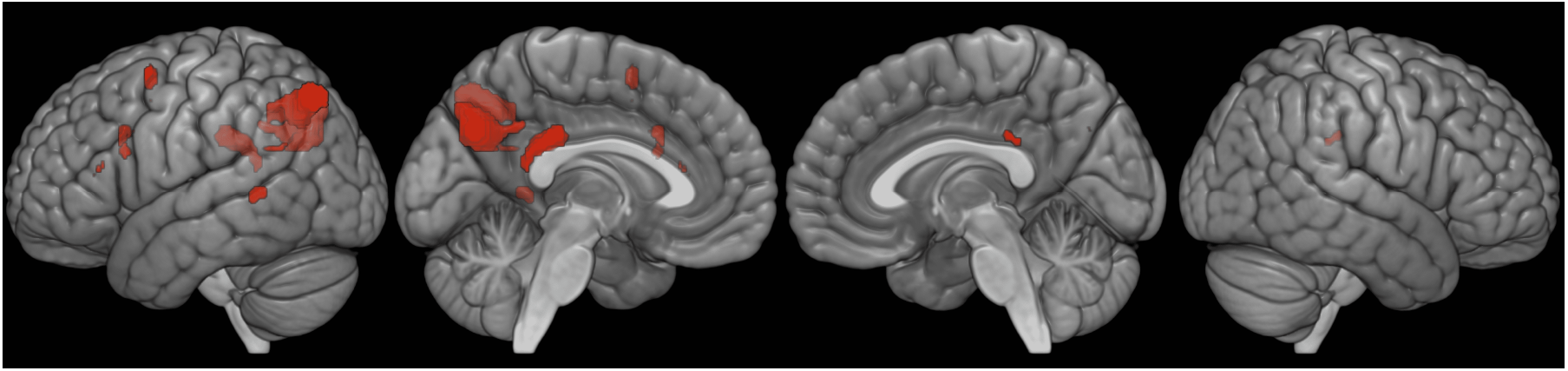
Object-extracted relative clause voxel-level intersection of the false discovery rate-thresholded Chinese and English maps, with no cluster thresholding.

**Table 4.** Object-*extracted* relative clause voxel-level intersection of the false discovery rate-thresholded English and Chinese results, with no cluster thresholding. Region, cluster size, and MNI coordinates.

| Region | Cluster size<br>(mm<sup>3</sup>) | MNI coordinates |  |  |
| --- | --- | --- | --- | --- |
|  |  | x | y | z |
| L Middle Temporal Gyrus (BA 21) | 304 | -60.0 | -44.0 | 4.0 |
| L Angular Gyrus (BA 39) | 96 | -52.0 | -54.0 | 24.0 |
| L Angular Gyrus (BA 39) | 3576 | -40.0 | -64.0 | 44.0 |
| L Pars Opercularis (BA 44) | 104 | -46.0 | 16.0 | 22.0 |
| L Pars Triangularis (BA 45) | 32 | -46.0 | 28.0 | 14.0 |
| L Pars Triangularis (BA 45) | 8 | -48.0 | 30.0 | 12.0 |
| L Pars Triangularis (BA 45) | 24 | -46.0 | 26.0 | 16.0 |
| L Premotor Cortex (BA 6) | 240 | -40.0 | 4.0 | 56.0 |
| L Frontal (BA8) | 240 | -36.0 | 16.0 | 30.0 |
| L Premotor Cortex (BA 6) | 8 | -36.0 | 4.0 | 46.0 |
| L Frontal (BA8) | 8 | -32.0 | 12.0 | 32.0 |
| L Precuneus (BA 7) | 3824 | -8.0 | -66.0 | 34.0 |
| L Posterior Cingulate Cortex (BA 23) | 1424 | -4.0 | -36.0 | 26.0 |

## Discussion

The central finding is one of commonality. Across Chinese and English, there was voxel-level overlap in the left pMTG, left AG, and left IFG, including both the pars opercularis and pars triangularis. This overlap extended to left premotor cortex, left precuneus, and posterior cingulate cortex. Such commonality bolsters the idea of a uniform brain basis for ORC processing. Further supporting this point of neural uniformity, there were no significant differences between the English- and Chinese-specific activations: at no location in the brain was ORC processing in one language associated with significantly greater activation than ORC processing in the other.

The regions identified under the rubric of this commonality subsume those which have previously been found for ORC processing in English: left IFG (Caplan et al., 2008; Just et al., 1996; Stromswold et al., 1996), right IFG (Just et al., 1996), left pM/STG (Caplan et al., 2008; Just et al., 1996), and right pM/STG (Just et al., 1996). They fit squarely into the traditional language network, and have also been found for ORC processing in Chinese: left IFG (Xiong & Newman, 2021), right IFG (Xiong & Newman, 2021), left pSTG (K. Xu & Duann, 2020; K. Xu et al., 2020b), right pSTG(K. Xu et al., 2020b), left mid M/STG (Lee & Chan, 2023), right mid M/STG (Lee & Chan, 2023), left MTG (Xiong & Newman, 2021; K. Xu & Duann, 2020; K. Xu et al., 2020b), right MTG (K. Xu et al., 2020b), left MTG broadly (Xiong & Newman, 2021), and right MTG broadly (Xiong & Newman, 2021).

Common areas extend beyond the traditional language network, viz left AG, left premotor cortex, left precuneus, and posterior cingulate cortex. They are evidently specific to ORCs. That is, they are not accounted for by any of GLM control predictors such as lexical meaning, non-relative clause syntactic structure, or word-string surprisal. This goes beyond certain classic findings based on non-naturalistic stimuli in English (although see Caplan et al., 2008, for a left precuneus result). The involvement of such extended areas is consistent with more recent neuroimaging results from Chinese, which identify left AG (Xiong & Newman, 2021), right AG (Xiong & Newman, 2021), left premotor cortex (K. Xu et al., 2020b), left precuneus (Xiong & Newman, 2021), and left posterior cingulate cortex (K. Xu et al., 2020b) for ORC processing.

The mismatch between classic results with English and the more recent Chinese results, including those reported here, may boil down to the choice of baselines. In the English neuroimaging studies discussed above, ORC effects arose either from an ORC-sentence *>* SRC contrast (Caplan et al., 2008), an ORC-sentence *>* sentence-with-a-single-nonword contrast (Stromswold et al., 1996), or from an region-of-interest analysis which only considered the traditional language network (Just et al., 1996). This is in contrast to most of the more modern Chinese results, where ORC effects are observed using a visual orientation baseline (K. Xu & Duann, 2020; K. Xu et al., 2020b), a fixation cross baseline (Xiong & Newman, 2021), or independent component analysis (Xiong & Newman, 2021). The only exception appears to be Lee and Chan (2023), who contrast ORC-sentence *>* sentence-baseline, and only identify the left IFG, bilateral MTG, bilateral STG, and right premotor cortex: essentially the traditional language network identified in the English ORC literature. When sentences are contrasted against one another (Caplan et al., 2008; Lee & Chan, 2023; Stromswold et al., 1996), effects within the extended language network seem to cancel each other out. Only when the ORC constructions are contrasted against a non-linguistic baseline are effects observed in the “extended” areas.

The “extended” areas observed in this study are all familiar from previous naturalistic studies of both English and Chinese: precuneus (Lerner et al., 2011; Maguire et al., 1999; J. Xu et al., 2005), posterior cingulate (Ferstl et al., 2008; Fletcher et al., 1995; Maguire et al., 1999), lateral frontal lobe (Lerner et al., 2011; Maguire et al., 1999; J. Xu et al., 2005), and angular gyrus (Lerner et al., 2011; J. Xu et al., 2005). Comparative naturalistic studies involving other languages, such as Farsi and Russian, is likewise consistent with the idea of an extended network being normally recruited in the service of ordinary, contextualized language comprehension (Dehghani et al., 2017; Honey et al., 2012).

These commonalities, within the traditional language network, are consistent with a variety of large-scale neurobiological models. The co-activation of both temporal and frontal areas would be explained, on the Extended Argument Dependency model, by appeal to the fact that these ORCs include both language-specific sequential aspect (prenominal vs postnominal) as well as filler-gap dependency that is independent of word order (Bornkessel-Schlesewsky & Schlesewsky, 2015, eADM). An alternative view holds that key syntactic aspects of the comprehension task, in ORCs, are carried out in IFG (L. Chen et al., 2023; Friederici, 2016). Yet another alternative localizes the bulk of sentence-level processing to temporal regions (Flick & Pylkkänen, 2020; Matar et al., 2021; Matchin & Hickok, 2020).

The precise nature of any cooperation between these perisylvian regions remains a matter of debate. An inviting possibility, consistent with the eADM, would be for semantic roles to be initially assigned by (computations that occur in) temporal regions. As suggested by Caplan et al. (2008), these tentative semantic role assignments might be “checked” by a process localized to IFG. In noncanonical structures such as the ORCs treated here, such roles may need to be reassigned. This is analogous to role-reassignment that may occur in German object-first sentences. Indeed previous neuroimaging with German object-first sentences has analogously implicated regions in the frontal lobe (Meyer et al., 2012), consistent with the relative clause results obtained here with English and Chinese.

In addition to commonalities, there were also differences. The results indicate that Chinese speakers recruit a larger number of brain regions. These areas included right hemisphere premotor cortex and mid/anterior STG, bilaterally. However, these Chinese-specific effects were not found to be statistically significant in the whole brain contrasts comparing Chinese and English. This Chinese-specific hemodynamic activity could reflect temporary ambiguity that is uniquely present in the Chinese stimulus. That is to say, in English relative clauses are generally cued by a function word such as “that” “who” or “which” (see e.g. Just & Carpenter, 1987, pages 139 and 142). They are not generally held to exercise the human sentence processing mechanism’s ambiguity-resolution ability to any significant extent. By contrast, Chinese relative clauses do attest as many as four temporary ambiguities, ambiguities that are not resolved until at least the relativizer *DE* (as shown in Figure 1 of Jäger et al., 2015; see also Bulut et al., 2018; Hsiao & Gibson, 2003; C.-J. C. Lin & Bever, 2011; Vasishth et al., 2013). The recruitment of additional brain areas in our Chinese participants would be consistent with a reanalysis processes that operates to resolve these ambiguities. One apportionment that appears to be consistent with prior work is phrase-structural reanalysis in bilateral STG (see Lee & Chan, 2023, for a related line of reasoning regarding activation differences between Chinese RCs, which are gap-filler constructions, and Chinese topicalization constructions, which are filler-gap constructions). The findings reported here do not uniquely identify a particular functional localization, nor distinguish between “repair” and “reparsing” (as discussed by Grodner et al., 2003). However, they do align with the differential degree of ambiguity across the two languages.

## Conclusion

Despite their superficial typological differences, the brain basis of object relative processing seems to be largely uniform across English and Chinese. An extended set of brain areas seem to support ORC comprehension in both languages, elaborating the earlier picture of English that arose from non-naturalistic stimuli. More broadly, this study demonstrates how automatic annotation techniques, combined with a specific typological feature, may be used to investigate the brain bases of language from a cross-linguistic perspective (Bornkessel-Schlesewsky & Schlesewsky, 2016; Kemmerer, 2016).

## Notes

### Competing Interest Statement

The authors have declared no competing interest.

### Summary of Updates

Revision corresponding to peer review.

