## Supplemental Material for "Neural correlates of object-extracted relative clause processing across English and Chinese"

Donald Dunagan[1], Miloš Stanojević[2], Maximin Coavoux[3], Shulin Zhang[1], Shohini Bhattasali[4], Jixing Li[5], Jonathan Brennan[6], and John Hale[1]

[1]University of Georgia
[2]DeepMind
[3]Université Grenoble Alpes
[4]University of Toronto Scarborough
[5]City University of Hong Kong
[6]University of Michigan

### Identifying Chinese Object-Extracted Relative Clauses in the Stimulus Text

The main text section entitled "Stimuli and storybook annotations" describes a procedure for annotating regions of expected comprehension difficulty in English using a combination of parsing, tree geometry, and cognitive theory viz. the HOLD hypothesis of Wanner and Maratsos (1978).

The procedure for Chinese follows the same principles but is slightly more complicated. In particular a different set of tree-geometric properties are applied to identify object-extracted relative clauses based on `benepar`'s output.

(8)
```
(CP << (IP <, NP < (VP  < VV !< NP)) [< DEC  | < (CP < DEC) ] )
$++ (NP (!<</时候/ !<< /情况/ !<< /现象/)) )
```

The expression in 8 is designed to locate object-extracted relative clauses that use the particle 的 "de." Under Chinese Treebank Bracketing Guidelines (Xue & Xia, 2000, §2.6.1) these constructions are analyzed as CPs, complementizer phrases, adjoined to noun phrases (NP). This adjunction relationship is expressed by the first instance of CP in 8 standing in the `$++` or left-sister relationship with NP (first instance of NP on the second line of the expression). The adjoined CP must dominate a word tagged DEC, either immediately so or via exactly one other, intermediate CP. DEC is that tag to be used when 的 "de" occurs as part of a relative clause (Xue et al., 2005, §3.2). The requirement that the adjoined CP dominate relative-clause "de" is expressed by the subexpression of 8 in square brackets. Within the CP, there must be an Inflectional Phrase. This sentence-like category is abbreviated IP; for further information please consult Stowell (1981, page 67) or the pedagogical presentation in Haegeman (1991, §3.2). This sentence-like category must have a noun phrase as its first daughter. This requirement, expressed with the constraint `IP <, NP` insists that any matches must show an overt subject. That is to say, the gap must not be in the subject position. An additional constraint on this sentence-like category is that its verb phrase must include a main verb `VV` that does not fall into one of the Chinese Treebank's special categories (e.g. as listed in Xue et al., 2005, Table 3). The matching verb phrase must not include a noun phrase daughter, enforced by `VP !< NP`. This constraint requires that the gap occur in direct object position, i.e. that it be an Object-Extracted relative clause, rather than some other kind. Finally, the NP must not be headed by 时候 "time", 情况 "circumstance" or 现象 "phenomenon." These would be used in Complement Clauses (see Patterson, 2020, §1.1).

We excluded two additional cases on the basis of hand-checking by a trained linguist. They are both Adjunct Relative Clauses:

(9)

| | | | | | | | | |
|---|---|---|---|---|---|---|---|---|
| 这 | 是 | 一 | 座 | 玫瑰 | 盛开 | 的 | 花园 | 。 |
| This | is | a | Classifier | rose | blossoming | DE | garden | . |

"This is a garden that has roses blossoming."

(10)

| | | | | | |
|---|---|---|---|---|---|
| 第二 | 列 | 灯火 | 通明 | 的 | 快车 |
| The second | Classifier | lights | well-lit | DE | express |

"The second express with all lights well-lit"

As explained in further detail by Patterson (2020) such Adjunct RCs have a gap, but the gap is not an argument of the embedded verb. These examples are excluded because, linguistically, they are not comparable to the ORCs treated in the English half of this study.

### Parser Accuracy Scores

The section "Stimuli and storybook annotations" states that the English text was parsed using the `mtgpy` parser, (Coavoux, 2021) while the Chinese text was parsed using the `benepar` parser (Kitaev et al., 2019). We, here, report the performance of these systems. On the test section of the English Penn Treebank, the `mtgpy` parser achieves an F-measure of 95 (see Jurafsky & Martin, 2022, §4.7 for an overview of F-measure). On the test section of the Chinese Treebank 5.1, the `benepar` parser achieves an F-measure of 91.75.
